# Are entirely virus-free CAR T cells as good as lentiviral transduced universal cells?

**DOI:** 10.64898/2026.04.30.721956

**Authors:** Oliver Gough, Christos Georgiadis, Roland Preece, Renuka Kadirkamanathan, Waseem Qasim

## Abstract

Chimeric Antigen Receptor (CAR) T cells are now established as therapies for some haematological malignancies. While lentiviral or γ-retroviral vectors are commonly used for CAR delivery due to their efficiency and stable integration, supply constraints have created bottlenecks to wider applications and access. Alternatively, genome editing tools such as CRISPR-Cas9 can insert CAR genes by homology-directed repair (HDR) into specific genomic loci. ‘Universal’ donor CAR-T cells devoid of endogenous TCRαβ after CRISPR-Cas9-mediated editing of the T cell receptor alpha (*TRAC*) locus are being investigated for more cost-effective, ‘off-the-shelf’ therapies. Targeting insertion of CARs into the *TRAC* locus places transcription under the control of native regulatory machinery while simultaneously disrupting endogenous TCRαβ, and this has been reported to reduce exhaustion and extend persistence in modelling studies using humanised mice.

We compared anti-CD20 CAR-T cells, generated with CAR inserts at either *TRAC* or *CD3ζ* loci using entirely virus-free manufacture, and universal CAR20-T cells generated using existing lentiviral procedures and CRISPR/Cas9 knockout. While non-viral cell yields were lower than lentiviral products cytotoxic function *in vitro* was comparable between groups. Studies in humanised murine models of leukaemia inhibition found non-viral CAR20-T cells were generally less efficacious than LV-CAR20 and exhibited more exhausted phenotypes. Non-viral approaches offer the prospect of sophisticated editing and precise CAR insertion but careful preclinical evaluation and well-designed clinical trials benchmarked against lentiviral approaches are recommended.

## Introduction

CAR-T cells are typically manufactured using integrating viral vectors, most notably γ-retroviral or lentiviral agents, and vector manufacture and release testing is a major constraint on wider adoption. These vectors preferentially insert into transcriptional start sites and active genes respectively, and integration sites have been widely mapped (1). Occasionally clonal dominance has been documented in patients, for example lentiviral insertion into the *TET2* gene was reported to account for 90% of circulating CAR19-T cells in a patient with CLL (2), and insertions into the *CBL* gene locus were linked to clonal expansions in another patient (3). Insertional effects may be important for expansion, function and persistence and related to therapeutic efficacy but also influence longer term transformation risks. While broadly considered safe, CAR-T cell therapies have recently been under scrutiny for possible genotoxicity and are being assessed in the context of underlying predispositions and elevated risks of secondary cancer in heavily pre-treated patients (4). A small minority of secondary T cell malignancies investigated to date have harboured detectable vector copies (5). Nonetheless, alternative approaches that avoid uncontrolled insertion of a transgene and vector elements – including promoter and enhancer sequences – would help alleviate safety concerns and should provide a more streamlined, reduced-cost, path to T cell engineering.

Genome editing tools can support precise insertion of CAR transgenes using homology-directed repair (HDR) (6). Targeted insertion of CAR into the endogenous T cell receptor alpha chain constant (*TRAC*) locus has been widely investigated, and some murine studies suggested that CAR expression under the transcriptional influence of the endogenous *TRAC* promoter were superior with improved function and persistence (7). A variety of nucleases, including Transcription Activator-like Effector Nucleases (TALENs) (8), meganucleases (9) and CRISPR-Cas platforms (6,10) have been combined with template delivery through non-integrating viruses (mainly Adeno-associated virus, AAV) or by electroporation of plasmid or synthetic DNA. Improved cell handling and manufacturing processes have been developed with improved efficiency, viability and cell yields of CAR-T cells manufactured without viral vectors and resulting in more precisely defined modifications.

We previously described manufacture of therapeutic CAR-T cells using lentiviral transduction in combination with CRISPR-Cas9 or base editing to disrupt TCRαβ and other genes to generate ‘universal’ donor CAR-T cells (11,12). Here, we investigated virus-free products, generated without lentiviral vectors (LV), and compared these directly against LV-generated cells using a CD20-specific CAR against B cell malignancies. Efficiency of integration into the *TRAC* or *CD3ζ* loci was initially investigated using alternative forms of double-stranded (ds) DNA or single-stranded (ss) DNA and we tested enhancement strategies based on ‘double-tap’ editing or inclusion of small molecule inhibitors of non-homologous end joining (NHEJ) to improve yields of CAR20^+^TCRαβ^-^T cells. Existing clinical lentiviral configurations were combined with TCRαβ genome editing steps as a benchmark for viral CAR-T cell productions, providing direct quality and yield assessments, and functional comparisons at the end of production both *in vitro* and in humanised mice experiments. Our findings suggest carefully designed human studies against lentiviral benchmarks will be required to assess the therapeutic efficacy of alternative manufacturing processes and elucidate further improvements.

## Methods

### Cell lines

HEK-293T cells (RRID: CVCL_0063) were cultured in Dulbecco’s Modified Eagle Medium (DMEM; Gibco, Thermo Fisher Scientific, Waltham MA, USA) and Daudi B cell Burkitt’s Lymphoma line (RRID: CVCL_0008) in Roswell Park Memorial Institute (RPMI) 1640 medium (Gibco), both with 10% Foetal Bovine Serum (FBS; Gibco) and 1% Penicillin-Streptomycin (Gibco). Daudi lines which expressed a Firefly Luciferase / GFP cassette (Daudi-FLG) were produced as previously described (11).

### Plasmids

Homology donor template plasmids and lentiviral transfer plasmids were designed in Snapgene (3.1.4) and synthesised by Geneart (Invitrogen, Thermo Fisher Scientific, Regensburg, Germany), and lentiviral packaging plasmids were produced by Plasmid Factory GmbH (Bielefeld, Germany).

### Synthetic DNA repair templates

Linear dsDNA templates were either amplified from plasmids by PCR (Supplementary table 1) or purchased to include 300bp homology arms (unless specified).

dsDNA was produced by Touchlight Genetics (Hampton, UK) (doggy-bone DNA (dbDNA)) or Genscript Biotech (Nanjing, China) (GenWand dsDNA) and ssDNA by Genscript Biotech (GenExact ssDNA).

### Lentiviral vector production

Vesicular stomatitis virus G envelope protein (VSV-G) pseudotyped lentiviral vectors were produced using a 3^rd^ generation packaging system and concentrated via ultracentrifugation. as previously described (12).

### Flow cytometry

Flow cytometry staining was carried out in FACS buffer (PBS + 3% FBS) with antibodies listed in Supplementary table 3. Samples were acquired on a FACSymphony A5 cell analyser (BD Biosciences) using FACSDiva software (BD Biosciences, San Jose, CA, USA). Analysis was carried out in FlowJo version 10.8.1.

### Primary cell isolation and culture

Peripheral blood mononuclear cells (MNCs) were isolated from whole blood using Ficoll-Paque PLUS (Cytiva, Marlborough, MA, USA) or obtained from steady-state leukapheresis volunteer donations (Antony Nolan, London UK).

MNCs were activated for 48 hrs with TransACT T cell stimulation reagent (Miltenyi Biotec, Bergisch Gladbach, Germany) and cultured in TexMACS medium (Miltenyi Biotec) + 3% human serum + 100U/mL IL-2 (Miltenyi Biotec).

### Production of Lentiviral CAR-T cells

MNCs were transduced 24hrs after TransACT activation with pCCL-CAR20 lentiviral vector at an MOI of 5. 48-72hrs post-transduction, cells were electroporated with HiFi Cas9 RNP targeting *TRAC* or *TRAC* plus *CD38* using a Lonza 4D nucleofector. sgRNA (Supplementary table 2) was sourced either from Synthego or Integrated DNA technologies. On day 5 cells were moved to G-Rex cell expansion vessels (Wilson Wolf) and expanded for 7 days, with IL-2 supplementation every 2-3 days. Residual TCRαβ-expressing cells were depleted by column processing (Miltenyi) and cryopreserved (11).

### Production of non-viral CAR-T cells

MNCs were activated for 48hrs and then electroporated with HiFi Cas9 RNP targeting either *TRAC* or *CD3ζ* alongside DNA homology template. Cells were moved into a G-Rex cell expansion vessel on day 5 and processed as above.

### ‘Double-tap’ editing and NHEJ inhibitors

In double-tap editing HiFi Cas9 was complexed with a pool of sgRNAs containing the main *TRAC* sgRNA and 3 secondary sgRNAs targeting major InDel products as previously described (13,14).

In experiments with the NHEJ inhibitor, 1µM AZD7648 (Selleckchem) was supplemented for 24hrs post-electroporation, before transfer to fresh TexMACS medium (Miltenyi Biotec) + 3% human serum + 100U/mL IL-2 (Miltenyi Biotec) media (15).

### Molecular confirmation of editing

Genomic DNA was extracted with a DNeasy Blood and Tissue DNA kit (Qiagen). PCR was carried out using Q5 polymerase (NEB, Ipswich, MA, USA) (primers: Supplementary table 1).

To assess on-target editing, desired PCR bands were gel extracted and purified (Qiagen QIAquick, Hilden, Germany). PCR products were sent for sanger sequencing (Eurofins Genomics, Ebersberg, Germany) and analysed using ICE analysis software (ice.synthego.com).

In/Out PCRs primers were designed across breakpoints to be selective for inserted transgenes. Presence of in/out PCR bands was assessed by gel electrophoresis. Positive PCRs were purified and sequenced as detailed above.

### Nanopore sequencing and analysis

Genomic DNA libraries were produced using Ligation Sequencing Kit V14 (Oxford Nanopore SQK-LSK114, Oxford, UK), including recommended QC steps, and sequenced on a PromethION flow cell and with adaptive sampling enabled to enrich for sites of interest (MinKNOW 24.06.10; Dorado 7.4.12 with High-accuracy model v4.3.0, 400 bps). Sequencing was carried out by the UCL Genomics core facility.

Alignment to the hg38 reference genome or custom TRAC-CAR20 reference was carried out using the wf-alignment workflow (github.com/epi2me-labs) on the UCL Myriad High Performance Computing Facility.

Analysis of editing was carried out using the custom python tool AnalyseLongHDR. Briefly, aligned reads at the *TRAC* editing site were first filtered to remove those arising from episomal template DNA and non-informative reads. Reads with no transgene overlap were then binned as non-HDR reads (wild-type / small InDels). Remaining reads were then validated by motif matching for transgene-specific sequences. Reads with additional large insertions (>1000bp) were considered potential concatemers and validated with concatemer-specific motif matching.

### ddPCR copy number assessment

Assessment of CAR20 transgene copy number was carried out by duplex ddPCR assay using primers (Thermo Fisher Scientific) and probes (Integrated DNA Technologies, Coralville, IA, USA) (Supplementary table 1) targeting both the CAR scFv (FAM) and human albumin (HEX) as a normalisation reference. Reactions were run with ddPCR supermix for probe, no dUTP (BioRad) and the QX200 AutoDG Droplet Digital PCR system (Bio-Rad Laboratories, Hercules, CA, USA) was utilised for droplet generation and sample acquisition. Analysis was carried out using Quantasoft (Bio-Rad) and CAR transgene copies were normalised to the internal albumin reference to calculate transgene copies per cell.

### ^51^Cr cytotoxicity assay

^51^Cr-labelled Daudi cells were incubated with CAR-T cell products or unedited control T cells across a range of Effector: Target (E: T) ratios (4hrs, 37^°^C). CAR expression was normalised to the lowest-expressing group by dilution with donor-matched TCRαβ^-^ knockout T cells. Supernatants were then harvested and incubated with OptiPhase HiSafe 3 (PerkinElmer, Shelton, CT, USA) scintillation fluid for 16hrs at RT. ^51^Cr scintillation signal was measured on a Wallac 1450 MicroBeta TriLux microplate scintillation counter. ^51^Cr release resulting from specific lysis in co-culture wells was calculated as follows: [(Release_co-culture_ – Release_spontaneous_) / (Release_Max_ – Release_spontaneous_)⍰*⍰100].

### Cytokine release assays

CAR-T cells, normalised for CAR expression as described for ^51^Cr assays, were incubated with Daudi cells in triplicate (1:1 culture, 16-24hrs, 37^°^C). Supernatant was harvested and assayed the Human Th1/Th2/Th17 Cytometric Bead Array (CBA) kit (BD Biosciences).

### *In vivo* tumour suppression assays

NOD/SCID/γc ^-/-^ (NSG) mice (RRID: IMSR_JAX:005557) were inoculated intravenously (i.v.) with 5×10^5^ CD20^+^ Daudi-FLG cells (Supplementary methods) and disease engraftment was confirmed on day 3. Mice then received either CAR-T cells (4×10^6^ CAR^+^ cells i.v.) or unedited controls (10×10^6^ total cells i.v.), alongside PBS-only vehicle controls, again by tail-vein injection on day 4. For reduced dose experiments, 2.5×10^6^ CAR^+^ cells total were used.

Disease levels were tracked by weekly bioluminescent imaging using an IVIS Lumina III In vivo Imaging System (PerkinElmer, live image version 4.5.5). Bone marrow aspirates were harvested for assessment of tumour burden and residual T cell phenotyping by flow cytometry.

### Statistical analysis and Graphing

All graphing and statistical analysis was performed in Graphpad Prism version 10.

For mouse bioluminescence comparisons, area-under-the-curve (AUC) was calculated for each group and compared via a one-way ANOVA with multiple comparisons by a post-hoc Tukey’s test.

Survival curves were compared by pairwise log-rank tests, with Bonferroni correction for multiple comparisons (Family-wise threshold p<0.05; p_Bonferroni_ = 0.0083).

## Results

### Comparison of lentiviral and non-viral CAR20^+^ TCRαβ ^-^ T cell production

A CD20-specific CAR (CAR20) with a *CD8TM-41BB-CD3ζ* configuration was expressed by a pCCL lentiviral vector (LV-CAR20) under the control of an internal human PGK promoter as a current therapeutic benchmark (Figure 1A). To generate ‘universal’ LV-CAR20 T cells, CRISPR-Cas9 editing for TCRαβ disruption was incorporated by adding an electroporation step to deliver ribonucleoproteins (RNP) comprising SpCas9 and for *TRAC*-specific sgRNA (Figure 1A). Alternatively, for non-viral delivery, DNA templates were designed for in-frame insertion of a self-cleaving P2A peptide and CAR20 transgene into *TRAC* (TRAC-CAR20) or *CD3ζ* (CD3ζ-CAR20) which placed CAR expression under the control of 5’ endogenous transcriptional machinery at both loci. For *TRAC*, the insert included a polyA tail distal to the CD3ζ domain of CAR20, whereas for *CD3ζ*-sited CAR20 the template was modified to capture endogenous CD3ζ elements in-frame (Figure 1B). Importantly, template insertion at either *TRAC* or *CD3ζ* sites ensured simultaneous disruption of TCRαβ/CD3 assembly by preventing expression of TCRα or CD3ζ, an important feature of universal CAR20^+^TCRαβ^-^ T cells. For entirely virus-free engineering, electroporation after 48 hours of T cell activation delivered similar SpCas9 RNP complexes supplemented with template DNA for HDR-mediated site-specific insertion of homology flanked CAR20. For all products, after expansion, magnetic-bead-mediated depletion reduced residual TCRαβ ^+^ T cells to below thresholds likely to cause graft versus host disease (GVHD).

**Figure 1:**
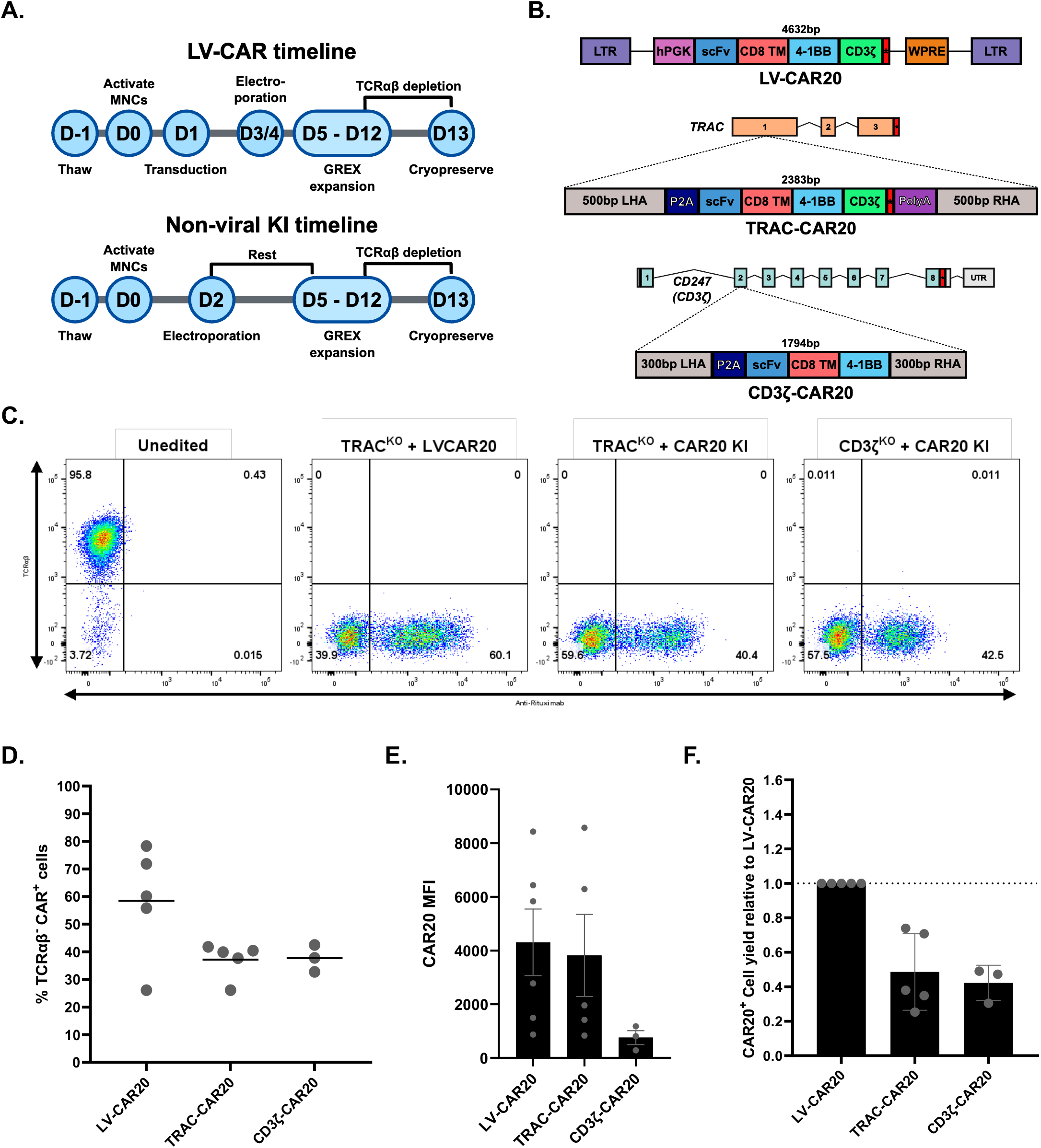
Generation of TRAC-CAR20 and CAR20-CD3ζ cells. **A**. Timelines for cell activation, transduction and editing for manufacture of LV or virus-free CAR20-T cells. **B**. Schematics showing the structure of the lentiviral CAR20 construct and CAR20 HDR templates targeting *TRAC* (TRAC-CAR20) and *CD3ζ* (CD3ζ-CAR20). **C**. Flow cytometry showing CAR20 expression and residual TCRαβ/CD3 at end of production (post-TCRαβ depletion) of lentiviral CAR20 cells with *TRAC* editing (LV-CAR20), and T cells with precise CAR20 insertion at *TRAC* (TRAC-CAR20) or *CD3ζ* (CD3ζ-CAR20). Non-edited T cells provided staining controls. **D**. Mononuclear cells (MNC) from multiple donors modified with LV-CAR20 and TRAC-CAR20 (n = 5) and CD3ζ-CAR20 (n = 3) showing TCRαβ^-^CAR^+^ fractions: LV-CAR20, mean 58.4% (*26*.*1-78*.*3%*); TRAC-CAR20, mean 37.1% (*26*.*1-41*.*8%*); CD3ζ-CAR20, mean 37.7% (*32*.*7-42*.*5%*). **E**. Mean Fluorescence Intensity (MFI) of CAR20 staining of TCRαβ-CAR^+^ fractions from CAR20 products. LV and TRAC-CAR20 were higher than for CD3ζ-CAR20. **F**. Final TCRab-CAR20+ numbers expressed relative to LV-CAR20 highlighted yields achieved with non-viral manufacturing were lower than for LV manufacturing. Each point represents one MNC donor, with mean and SEM.

Investigations compared different template DNA formulations. As expected, electroporation of standard bacteria propagated plasmid DNA compromised viability and CAR20^+^ TCRαβ^-^ T cell yields were poor (Supplementary figure S1A, B). Alternative DNA sources including PCR-amplified dsDNA, fermentation-produced covalently-closed dsDNA and synthetic covalently-closed linearised doggybone (db) DNA were investigated (Supplementary figure S1A, B) with the latter supporting 37.1% TRAC-CAR20 (26.1-41.8%, n = 5) and 37.7% CD3ζ-CAR20 insertion (32.7-42.5%, n = 3) (Figure 1C, D). For comparison, LV-CAR20 cells exhibited 58% CAR20 expression (26.1-78.3%, n = 5). LV-CAR20 and TRAC-CAR20 cells exhibited similarly broad and overlapping fluorescence intensities of CAR20 expression, whereas CD3ζ-CAR20 exhibited narrower, less intense mean fluorescence intensity (MFI) (Figure 1E). As anticipated CAR20 transduction and overall cell yields at the end of lentiviral production were higher for LV than for virus-free products with TRAC-CAR20: on average 52% lower and CD3ζ-CAR20 on average 58% lower (Figure 1F). Potential enhancement strategies included ssDNA templates, which supported improved cell viability but were less efficient (Supplementary figure 1C, D) and the introduction of additional sgRNAs for ‘double-tap’ editing (using sgRNAs against predicted InDel repair products) did not improve outcomes (Supplementary figure 2A). Addition of a small molecule NHEJ inhibitor AZD7648 increased HDR-mediated insertion (Supplementary figure 2B-D), but at the expense of viability and modified CAR20^+^ T cell yields were not improved (Supplementary figure 2E, F).

### Molecular signatures of site-specific CAR20 integration

Signatures of non-homologous end joining (NHEJ) effects following CRISPR-Cas9 activity at the *TRAC* and *CD3ζ* loci were confirmed by sequencing samples exposed to RNP but devoid of template HDR, with 88-92% indel formation documented in all three groups (Supplementary figure 3). Copy number assessments using ddPCR-detected background signals from non-integrated DNA template at early time points, necessitating additional control measurements after extended culture for over 21 days and non-edited template-only comparisons devoid of Cas9 or control sgRNA targeting *B2M* rather than *TRAC* (Supplementary figure 4). Quantification above background found 1.37 copies/cell for TRAC-CAR20 and 1.24 copies/cell for CD3ζ-CAR20, while LV-CAR20 cells harboured 2.59 copies/cell, comparable to clinical products (Figure 2A). To confirm correctly orientated insertion of CAR transgenes after HDR of DNA templates, targeted 5’ and 3’ PCR was performed at chromosomal breakpoints (Figure 2B; Supplementary figure 4) and amplicons were sequenced to verify in-frame transgene insertion. To further characterise template insertion, Nanopore whole genome sequencing (WGS) with adaptive sampling was used to enrich *TRAC* insertion sites. Alignment of reads to a putative reference sequence comprising the TRAC site with a CAR20 template inserted was complicated by detection of non-integrated template DNA which included flanking homology regions (Supplementary figure 6A, B). Reads filtered for high fidelity CAR integrants at the *TRAC* locus accounted for 43% of reads (Figure 2C; Supplementary figure 6C), with around 3.4% of reads harbouring larger insertions (>1Kb longer than anticipated) including concatemerisation events and other aberrant insertions.

**Figure 2:**
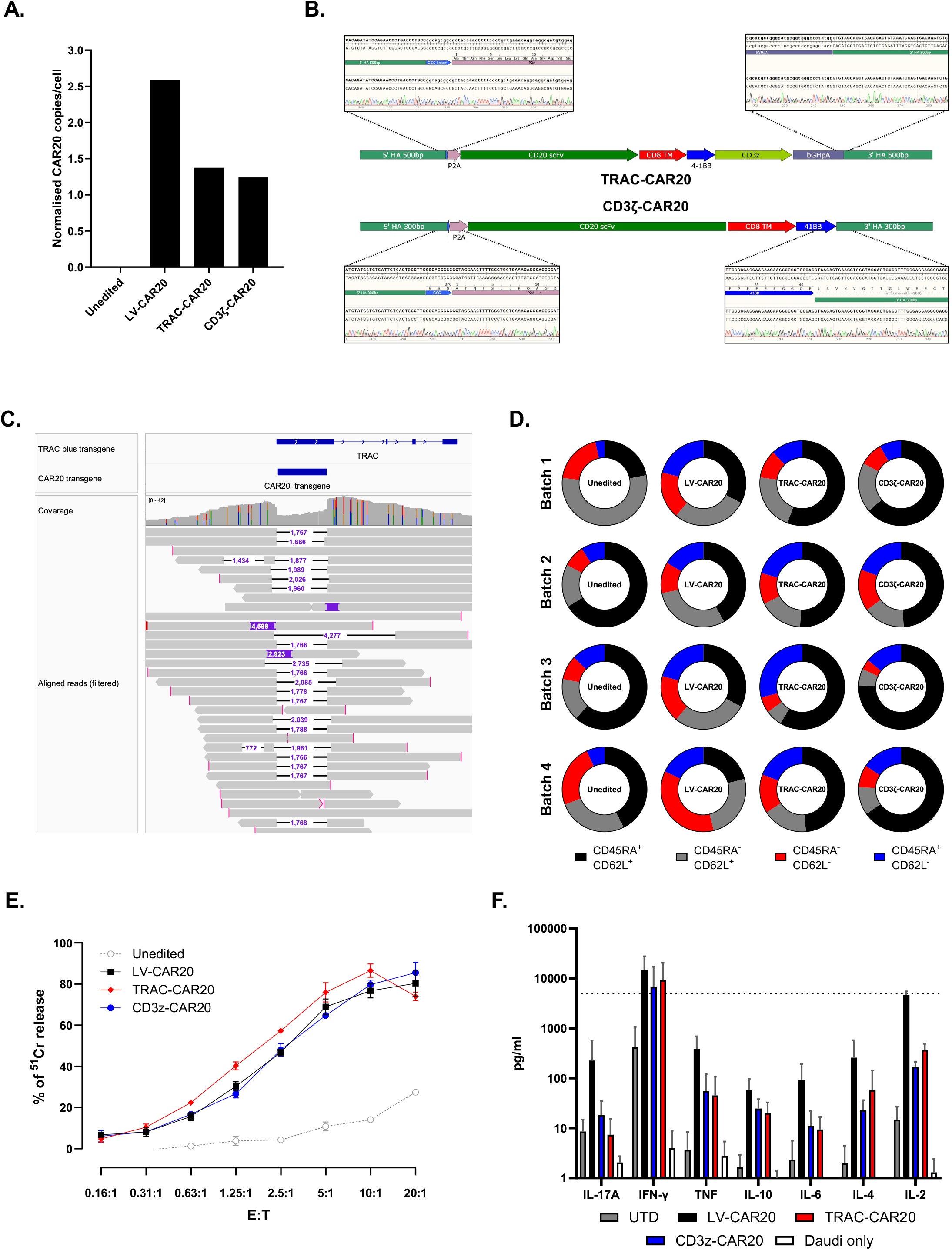
In vitro assessments of CAR20-T cell editing and function. **A**. Quantification of CAR20 transgene copies in each CAR-T cell product (Unedited, LV-CAR20: n=3; TRAC-CAR20, CD3ζ-CAR20: n=4). For TRAC-CAR20 and CD3ζ-CAR20 background readings (1.6 copies/cell) were attributed to non-integrated DNA. **B**. Schematic images of TRAC-CAR20 and CD3ζ-CAR20 with alignments of in / out PCR products showing detection of correctly inserted templates, with all expected breakpoints detected by electrophoresis (Supplementary figure 5). **C**. Additional long read nanopore sequencing & IGV alignment confirmed CAR20 transgene insertion into *TRAC* exon 1. Reads were filtered to remove suspected non-integrated template reads prior to analysis and visualisation. **D**. T cell phenotypes of unedited, LV-CAR20, TRAC-CAR20 and CD3ζ-CAR20 cells at end of manufacture, based on CD45RA and CD62L expression in CD2^+^ T cells, with no major differences between products (4 biological replicates; 1 per row). **E**. Cytotoxicity against Daudi target cells exhibited comparable antigen-specific killing by all CAR20 products. CAR-T cells from each group were co-cultured with ^51^Cr-loaded Daudi cells across a range of Effector: Target (E: T) ratios (Mean, SEM) **F**. Cytokine release in response to Daudi target, measured by cytometric bead array (CBA) assay after overnight coculture and harvest of supernatant (Mean, SEM; n=3), was also similar for all groups. Lines show upper and lower limits of quantitation.

### Virus-free and LV CAR20^+^ TCRαβ^-^ T cell characterisation in vitro

The phenotype of CAR20-T cell was characterised by flow cytometry for subsets including CD4/CD8 and effector-memory profiles based on surface expression of CD45RA and CD62L (T_EM_ : CD45RA-CD62L-; T_EMRA_ : CD45RA^+^CD62L^-^) (Figure 2D; Supplementary figure 7). After being normalised for CAR expression, all groups mediated comparable antigen-specific responses against CD20^+^ Daudi Burkitt’s lymphoma cells, both in terms of ^51^Cr cytotoxicity across a range of effector:target ratios (Figure 2E; Supplementary figure 8), and cytokine release profiles after overnight co-culture (Figure 2F). While end-of-production yields of CAR20^+^TCRαβ^-^ T cells were greater after LV transduction, there were no major differences in phenotype or function *in vitro* after normalising for CAR20 expression.

### Comparison of viral and non-viral CAR^+^ TCRαβ^-^T cells in humanised mice

CAR20^+^TCRαβ^-^ T cells were compared *in vivo* in a leukaemia inhibition model using NOD/SCID/γc^-/-^(NSG) immunodeficient mice pre-injected with Daudi tumour cells expressing luciferase & GFP (Daudi-FLG cells). Weekly bioluminescence imaging (BLI) assessed tumour burden (Figure 3A, B) and survival (Figure 3C) for up to 8 weeks. Over the first 2-3 weeks, leukaemia inhibition was comparable across effector groups with no significant differences between viral and non-viral products based on bioluminescence intensity (Figure 3B; Supplementary figure 9). Over the subsequent four weeks, disease progression in LV-CAR20 mice was impeded compared to non-viral CAR20 groups (Figure 3C). Mice survived significantly longer after LV-CAR20 cells compared to TRAC-CAR20 cells (p<0.005) or CD3ζ-CAR20 cells (p< 0.0001). Bone marrow samples recovered at necroscopy found TRAC-CAR20 and CD3ζ-CAR20 groups had readily detectable residual leukaemia cells, while tumour counts in LV-CAR20 treated mice were below the limit of quantification (Figure 3D). CD2^+^ CAR20^+^ T cell populations were detected in all groups with CD8^+^ dominance in TRAC-CAR20 and CD3ζ-CAR20 treated groups (Figure 5E; Supplementary figure 10B) although these cells also exhibited higher levels of exhaustion markers, PD-1, TIM3 and LAG3 (Figure 2F). Overall, in three direct independent comparisons, leukaemia inhibition and animal survival mediated by TRAC-CAR20 or CD3ζ-CAR20 T cells was inferior or at best comparable to control provided by LV-CAR20 T cells. (Figure 3; Supplemental figure 9 and 11)

**Figure 3:**
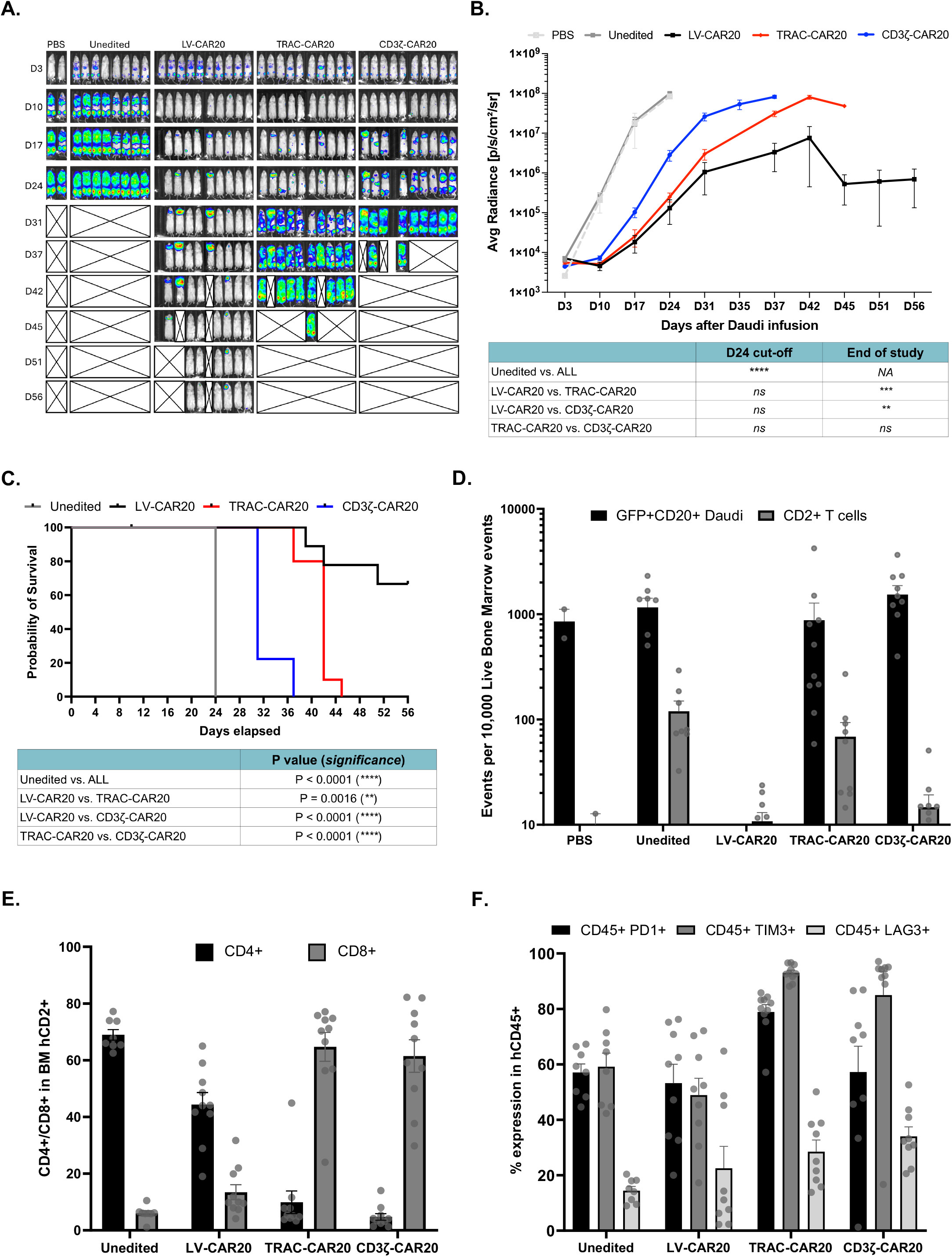
In vivo comparison of LV and non-viral CAR20 cells. **A**. Bioluminescent imaging (BLI) of NOD/SCID/γc^-/-^(NSG) mice engrafted with Daudi-FLG cells and dosed with either LV-CAR20, TRAC-CAR20 or CD3ζ-CAR20. Animals were followed for up to 56 days after tumour engraftment. Mice receiving non-modified T cells or PBS alone provided controls. **B**. Mean radiance values from BLI of Daudi-FLG leukaemia progression in NSG murine models (PBS control, n = 2; Unedited control, n = 8; for all other groups n = 10). By 56 days, LV-CAR20 exhibited significantly improved tumour control. Statistical analysis was carried out by one-way ANOVA of the area-under-the-curve (AUC) for each group, followed by a post-hoc Tukey’s test for individual comparisons. **C**. Overall survival was longer for LV-CAR20 than non-viral groups when Kaplan-Meier curves were compared (pairwise log-rank tests to a Bonferroni-corrected threshold of p<0.05). **D**. Daudi-FLG cell counts and CD2^+^ T cell counts in bone marrow (BM) at necroscopy highlighted residual GFP^+^ Daudi signal in non-viral groups but not LV-CAR20. Cells were measured by flow cytometry normalised to events per 1×10^5^ live cell events (mean with SEM) (Unedited, n = 8; LV-CAR20, n = 9; TRAC-CAR20, n = 10; CD3ζ-CAR20, n = 9). **E**. CD4^+^ and CD8^+^ T cell fractions in CD2^+^ cells recovered from the bone marrow of mice at point of necroscopy showed expanded CD8^+^ populations in non-viral groups (Unedited, n = 8; LV-CAR20, n = 9; TRAC-CAR20, n = 10; CD3ζ-CAR20, n = 9). **F**. Expression of PD-1, TIM3 and LAG3 in CD45^+^ cells recovered from the bone marrow showed increased exhaustion in non-viral groups (Unedited, n = 8; LV-CAR20, n = 9; TRAC-CAR20, n = 10; CD3ζ-CAR20, n = 9). P value summaries: p≥0.05 (ns), p<0.05 (^*^), p<0.01 (^**^), p<0.001 (^***^), p≤0.0001 (^****^).

Additional investigations tested 1:1 co-infusion of LV-CAR20 + TRAC CAR20 to investigate if in vivo competition effects arose for the two products over a 10-week period using serial BLI of Daudi disease burden, compared to each product alone. Disease course and survival was superior in CAR20 groups compared to controls, but there were no significant differences between effectors used alone or when co-infused (Supplemental figure 11). Residual leukaemia burden (Daudi-GFP) detection by flow cytometry of bone marrow (BM) at necroscopy found that animals dosed with TRAC-CAR20 alone had significantly higher residual counts compared to animals receiving LV-CAR20 alone or with TRAC-CAR20 (p<0.01). CD2+ human T cells were detectable at low levels by flow in all groups, but CAR20 transgene detection by CAR20 specific ddPCR, found signals were highest in LV-CAR20 animals and barely detectable in TRAC-CAR20 indicating reduced persistence of non-viral CAR T cells by the end of the experiment. Analysis of exhaustion marker expression suggested similar profiles across CAR20 groups, albeit across limited cell numbers, and elevated PD1 in controls animals which received non-edited TCRαβ+ T cells capable of xeno-reactive responses.

## Discussion

The fidelity and efficiency of precise nuclease-mediated DNA scission and template insertion by HDR have improved considerably in recent years. Initial approaches for CAR gene insertion in T cells employed Adeno associated virus (AAV) for episomal delivery of homology-flanked DNA repair templates, and supported CAR insertion into the TRAC site, safe harbour or checkpoint pathway loci (7,16). Clinical trials using such approaches have been underway, but there has been limited reporting of outcomes to date (17,18). The strategy has needed high AAV multiplicity of infection resulting in the same attendant issues of virus production and supply constraints that have affected LV approaches. Alternative ‘virus-free’ template delivery using different forms of DNA, including plasmids, minicircles, PCR-amplified dsDNA or synthetic ssDNA have also been investigated (6,19-22), with some approaches reaching clinical stage testing. For example, a recent trial for relapsed non-Hodgkins lymphoma described using dsDNA donor templates derived from linearized plasmids for HDR insertion of CAR genes, and another used plasmid delivery of neoantigen-specific TCRs to the *TRAC* locus for patients with refractory solid cancers (16,23). We compared in-house plasmid DNA with alternative sources of template DNA, including amplified PCR products, synthetic dsDNA and ssDNA. As anticipated we found that manufacture of CAR20 cells using bacteria-derived plasmid DNA resulted in poor viability and low efficiency of HDR, likely a consequence of innate DNA sensing in T cells (24,25). It has been reported that use of ssDNA templates can obviate effects of innate sensing of dsDNA while supporting high efficiency gene transfer (19), but we found editing efficiency was reduced. More favourable engineering was achieved using synthetic linear dsDNA templates which supported higher editing efficacy and supported greater CAR T cell yields at the end of production, albeit still notably lower than after standard lentiviral manufacturing.

Linearised synthetic dsDNA templates supported the most efficient insertions and end-yields. These were commercially manufactured using bacteria-free rolling circle amplification and enzymatic processing, and were devoid of bacterial components and associated epigenetic marking (26,27),that may otherwise be highly immunogenic (28).

A number of enhancement strategies to improve HDR-mediated transgene integration have been reported. Small molecule inhibitors of NHEJ or mismatched end joining) have been used to skew cell repair pathways after dsDNA breakage towards HDR rather than NHEJ (29,30). We found that the NHEJ inhibitor AZD7648 increased transgene integration, but final cell yields were reduced due to toxicity, consistent with concerns reported by others (31). Alternatively, ‘double-tapping’ through inclusion of additional sgRNAs targeting primary InDel products predicted to arise from TRAC sgRNA activity with the aim of secondary editing and additional HDR mediated CAR insertion (13,14). Such “recursive” editing did not improve HDR efficiency in our experiments and given that there may be additional off-target concerns the approach was not pursued further.

Overall, in vitro functional studies, when normalised for CAR expression, did not detect differences between non-viral CAR20 and LV-CAR20 cells. However, during *in vivo* T cell comparisons, disease in non-treated animals progressed more rapidly than CAR20 groups with divergence between LV-CAR20 and non-viral CAR20 groups after around three weeks in most groups. Previously, Eyquem et al. had suggested CAR expression under the control of

*TRAC* endogenous promoter machinery was superior to LV-CAR in a ‘stress test’ animal model (7). Rather than a dose-titrated stress model, our experiments employed an *in vivo* potency model that we have previously reported and used for testing LV-CAR T products ahead of clinical trials. We reasoned that any comparator product should perform at least as well as the current LV-CAR platform after normalising for CAR expression and we found that this benchmark was not surpassed by non-viral CAR20 products in any of our in vivo modelling. Others have reported comparisons of LV or CRISPR-Cas knock-in of CAR19 at similar TRAC sites and had suggested comparable performance in humanised mouse experiments, although these were undertaken over shorter periods and may not have captured later divergening outcomes between the groups (17). It is plausible that CAR insertion into other loci may confer additional advantages. For example, CAR integration into sites such as *PDCD*1 may simultaneously modify checkpoint pathways and tackle exhaustion (16). Other sites may support clonal dominance as reported after LV-CAR integration into sites such as *TET2* (2,32,33) or *CBL* (34). Deliberately targeting CAR insertion to sites such may drive expansion, although both *TET2* and *CBL* loss-of-function changes have been implicated in haematological malignancies (35,36). Nonetheless careful screening for integration sites which offer favourable CAR expression kinetics and promote persistence without creating adverse risks may improve CAR-T cell products.

In summary, non-viral manufacturing of ‘universal’ CAR20-T cells using CRISPR-Cas9 and synthetic DNA templates delivered precise transgene integration and simultaneous TCRαβ disruption, albeit with yields less than for LV-CAR20 manufacture. Whilst function of these cell products was similar to LV-CAR20 cells *in vitro*, the latter generally mediated more efficacious leukaemia clearance in vivo. Mimicking LV-mediated integration effects for more sophisticated recapitulations of vector-mediated variegation effects might be recruited for beneficial impacts on expansion and persistence, but in the meantime human studies will need to be carefully designed to allow comparisons against existing viral benchmarks.

## Acknowledgments

Supported by Medical Research Council, Wellcome Trust, National Institute of Health Research. Anthony Nolan Trust (803716/SC038827) supplied MNC donations. The authors acknowledge UCL Genomics and UCL Flow Cytometry core facilities, UCL Myriad High Performance Computing Facility; Ailsa Greppi, Kyle O’Sullivan supported *in vivo* studies; Hala Aldahshan designed site-specific PCR assays.

## Data availability

Sequencing data will be available via request to authors and code for analysis of HDR reads from nanopore sequencing is available at github.com/DrOJG/AnalyseLongHDR.

