## Supplemental Figures and Tables for "Are entirely virus-free CAR T cells as good as lentiviral transduced universal cells?"

### Supplementary figures and tables

**Supplementary figure 1:** Comparison of homology donor template options. *p.2*

**Supplementary figure 2:** Potential strategies to increase transgene insertion rates. *p.4*

**Supplementary figure 3:** ICE analysis of editing at *TRAC* and *CD3ζ*
in end-of-production CAR20 cells. *p.5*

**Supplementary figure 4**: CAR20 transgene copy number assessment. *p.7*

**Supplementary figure 5:** In / out PCR analysis gels for detection of
TRAC-CAR20 and CD3ζ-CAR20 insertions. *p.8*

**Supplementary figure 6:** Additional nanopore analysis figures. *p.9*

**Supplementary figure 7:** CD4 / CD8 expression in the $CAR20^{+}$
subpopulation at end-of-production. *p.11*

**Supplementary figure 8:** Additional donor replicates for $^{51}Cr$ cytotoxicity assays**.** *p.12*

**Supplementary figure 9:** Additional human murine chimeric data
comparing LV-CAR20, TRAC-CAR20 and CD3ζ-CAR20. *p.13*

**Supplementary figure 10:** Additional phenotyping of CAR-T cells recovered
from main *in vivo* comparison of LV-CAR20, TRAC-CAR20 and CD3ζ-CAR20. *p.14*

**Supplementary figure 11:** Single versus competitive LV-CAR20 / TRAC-CAR20 infusion. *p.15*

**Supplementary table 1:** PCR primers and ddPCR probes. *p.17*

**Supplementary table 2:** sgRNA sequences. *p.18*

**Supplementary table 3:** Antibodies used in study. *p.19*

**Supplementary figure 1: Comparison of homology donor templates**

Comparison of $CAR20^{+}$ $TCR\alpha\beta^{-}$ production using DNA templates encoding homology flanked-CAR20 sequences supplied as plasmid (Qiagen) or generated by bacteria-free synthetic production as single stranded DNA (Genscript, China), or double stranded DNA options dbDNA (Touchlight, London, UK) or dsDNA (Genscript, China).

**A.** Cell viability measured by flow cytometry (BD Horizon Fixable Viability Stain 700) for TRAC-CAR20 and CD3ζ-CAR20 cells made with plasmid DNA had substantially lower viability than cells generated using dbDNA.

**B.** Flow cytometry plot from a comparison of generation of CD3ζ-CAR20 cells using plasmid DNA and dbDNA. Plasmid DNA did not produce satisfactory CAR20 insertion.

**C.** Cell viability of TRAC-CAR20 cells generated with dbDNA, dsDNA or ssDNA templates at different time points across culture period.

**D.** TRAC-CAR20 cell production using dbDNA or dsDNA templates was superior to ssDNA template products in terms of KI efficiency.

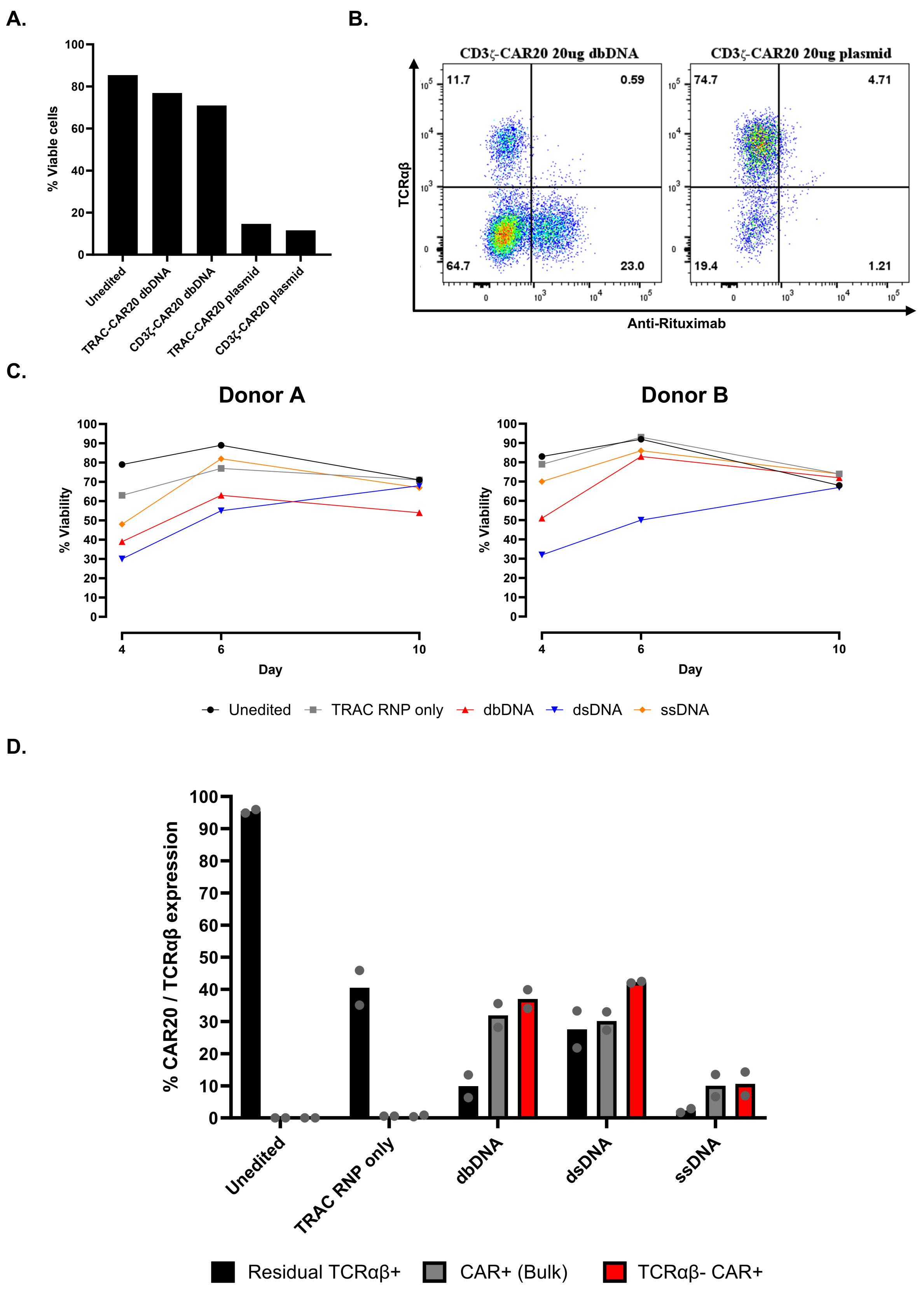

A.
100
80
60
40
20
0
% Viable cells
Unedited
TRAC-CAR20 dbDNA
CD3ζ-CAR20 dbDNA
TRAC-CAR20 plasmid
CD3ζ-CAR20 plasmid
B.
CD3ζ-CAR20 20ug dbDNA
CD3ζ-CAR20 20ug plasmid
TCRαβ
Anti-Rituximab
11.7
0.59
64.7
23.0
74.7
4.71
19.4
1.21
C.
Donor A
Donor B
% Viability
Day
4
6
10
Unedited
TRAC RNP only
dbDNA
dsDNA
ssDNA
D.
% CAR20 / TCRαβ expression
Unedited
TRAC RNP only
dbDNA
dsDNA
ssDNA
Residual TCRαβ+
CAR+ (Bulk)
TCRαβ- CAR+

**Supplementary figure 2: Potential strategies to increase transgene insertion rates.**

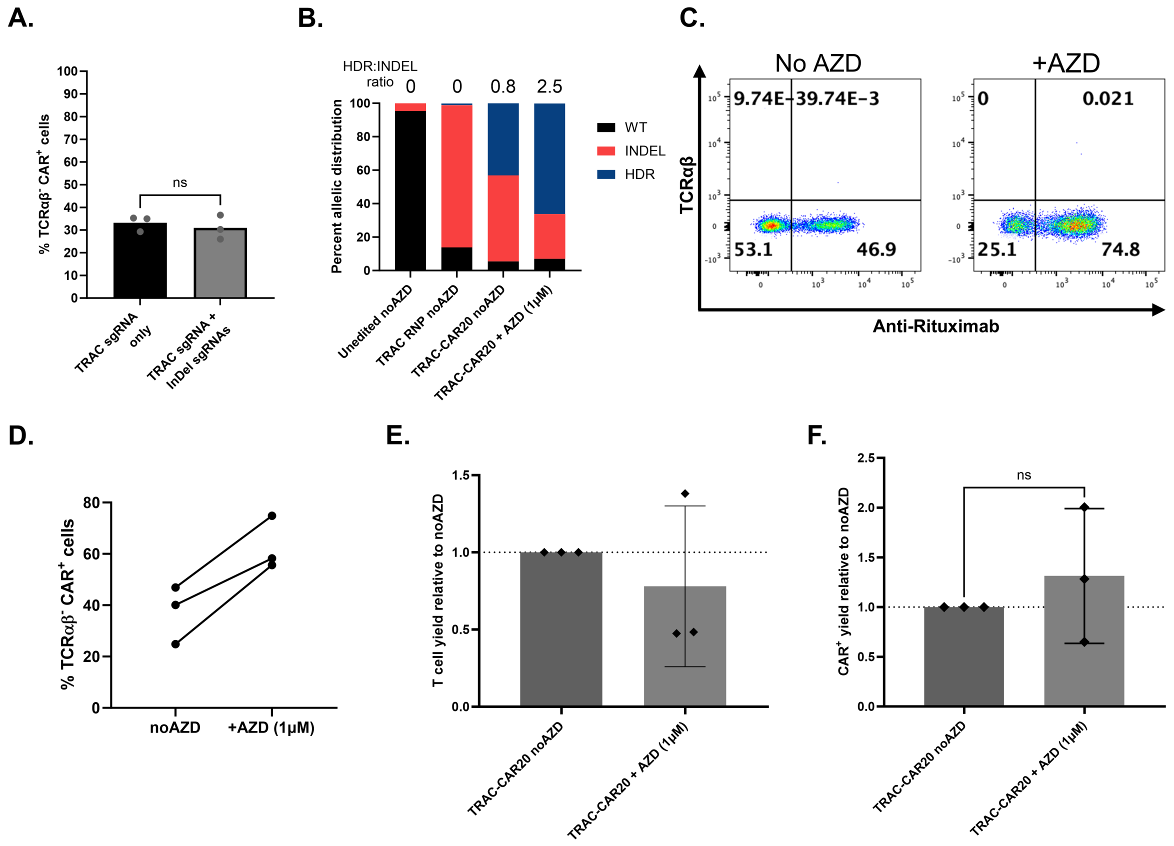


Strategies to increase site-specific CAR insertion by HDR included 'Double-tap' editing – whereby additional sgRNAs were used to target predicted InDel repair products of initial sgRNAs in order to re-edit for further HDR occurring but did not increase the rate of site-specific insertion (**A**). Manipulation of DNA repair pathways using NHEJ inhibition with the small molecule AZD7648 (AZD) increased CAR insertion, but reduced overall cell yields, therefore CAR$^+$ cell yields were not significantly improved.

**B.** Pre-depletion flow cytometry data was used to estimate the fractions of WT, InDel and HDR alleles present in the edited cell pool. Illustrated by the HDR:InDel ratios, addition of 1µM AZD7648 increased the fraction of HDR events while reducing the fraction of InDel events.

**C.** Representative flow cytometry plots showing TRAC-CAR20 cells made with or without AZD7648 after TCRαβ depletion, illustrating increased CAR20 insertion with the drug.

**D.** Summary graph showing the increase in the percentage TCRαβ$^-$CAR$^+$ cells with and without AZD7648 treatment. AZD7648 consistently increased the fraction of TCRαβ$^-$CAR$^+$ cells. n = 3.

**E.** Total T cell yields normalised to yield without AZD7648. Inclusion of AZD7648 generally led to reduced T cell yields. n=3.

**F.** CAR$^+$ cell yields post-depletion, normalised relative to TRAC-CAR20 cells made without AZD7648 treatment. While AZD7648 on average increased the yield of CAR$^+$ cells, this effect was inconsistent and non-significant, and CAR$^+$ yields were still substantially below those observed with lentiviral transduction. n = 3. Statistical comparison was carried out using an unpaired students T test.

**Supplementary figure 3: ICE analysis of editing at *TRAC* and *CD3ζ* in end-of-production CAR20 cells.**

**A.**

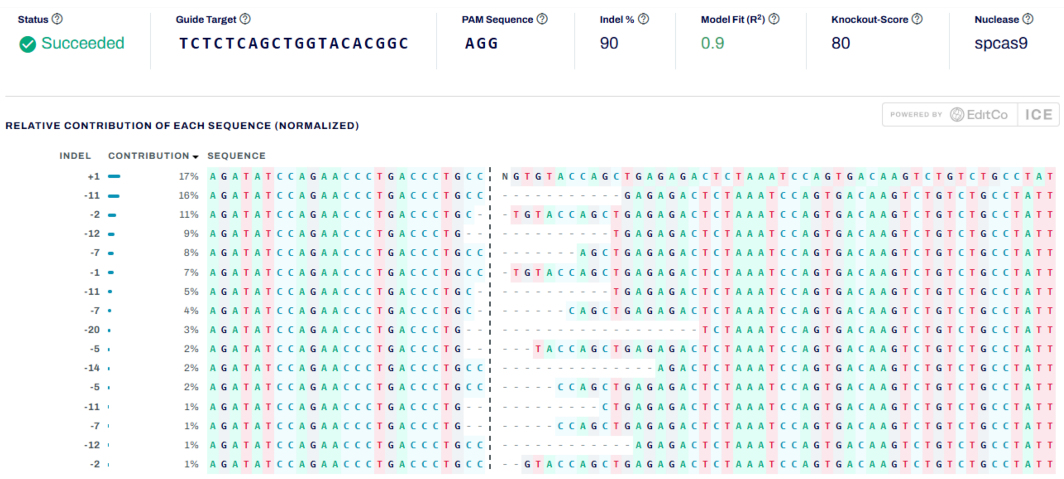


**B.**

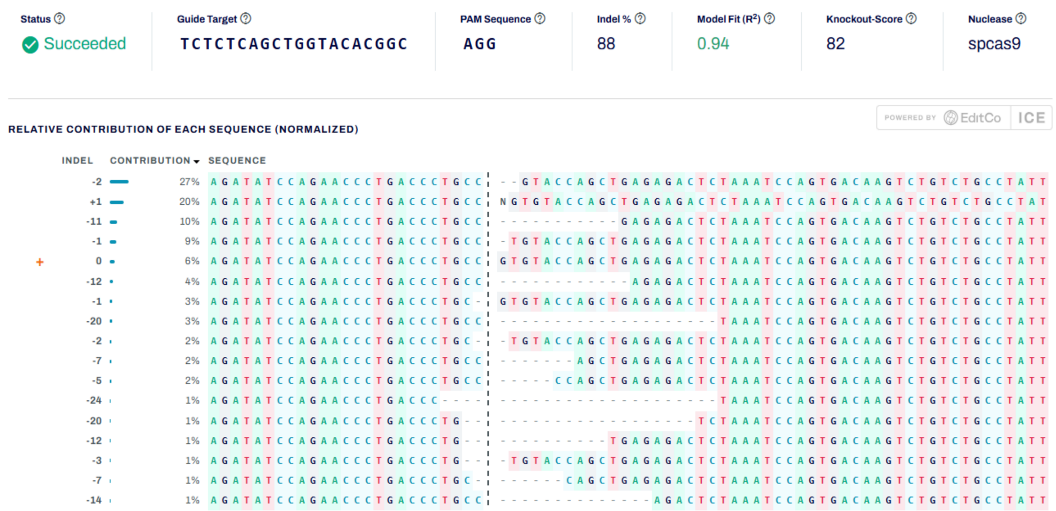


**C.**

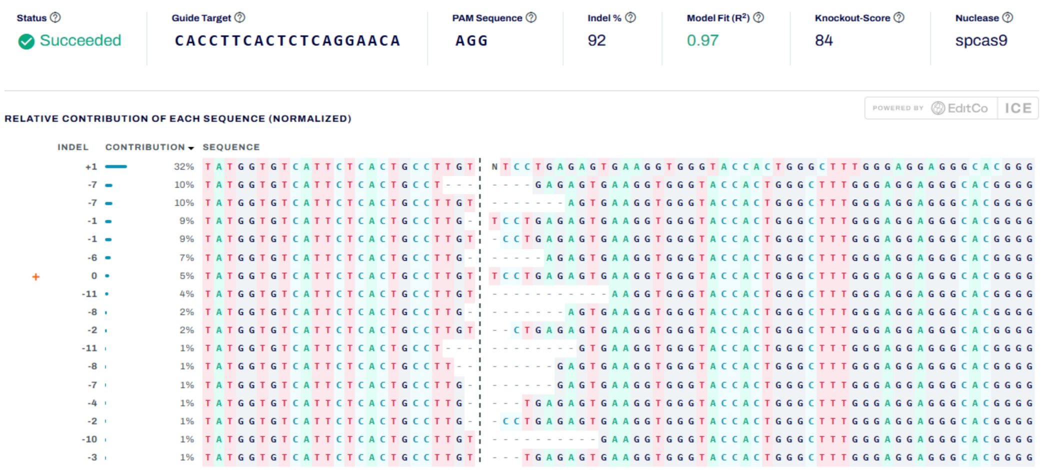

**Supplementary figure 3 (cont.):**

Representative analysis of InDel profiles from sanger sequencing traces using Synthego's ICE tool. Analyses shown are from **A.** the *TRAC* editing site in LV-CAR20 cells, **B.** The *TRAC* editing site in the non-HDR fraction of TRAC-CAR20 cells, **C.** the *CD3ζ* editing site in the non-HDR fraction of CD3ζ-CAR20 cells.

**Supplementary figure 4**: **CAR20 transgene copy number assessment**

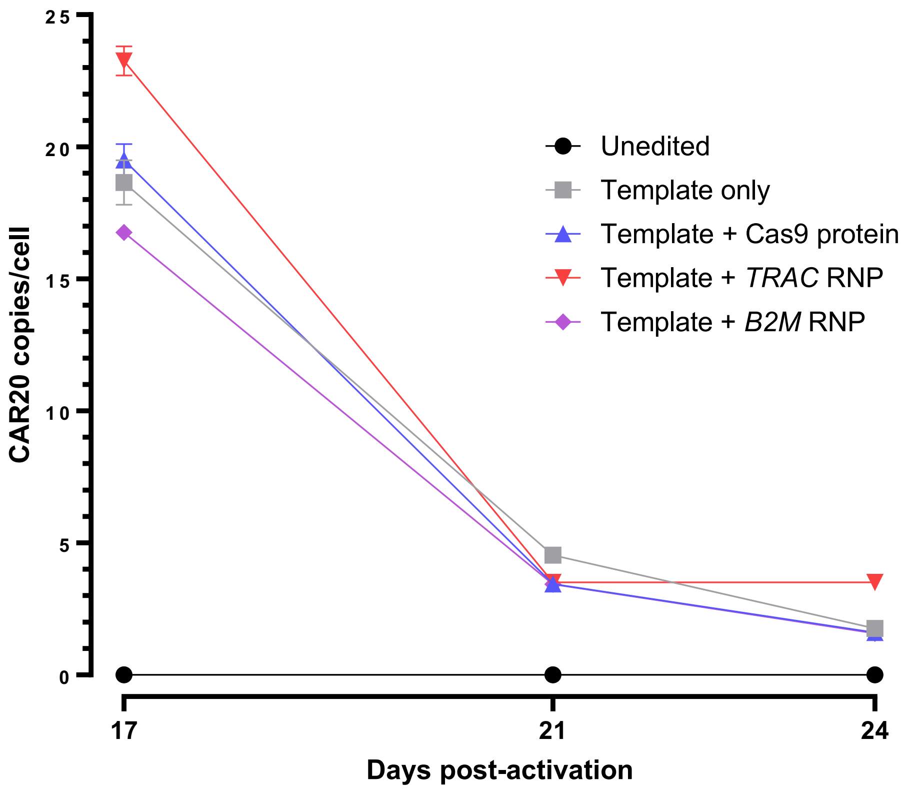


Time course experiment of primary T cells electroporated with either TRAC-CAR20 dbDNA template alone (Template only), dbDNA with Cas9 protein without an sgRNA (Template + Cas9), dbDNA with a Cas9 RNP targeting the *TRAC* locus, or dbDNA with a Cas9 RNP targeting the *B2M* gene locus. At 24 days, the *TRAC* KI copy number approximated that of End-of-Production CAR20 KI cells, giving an average background value of 1.6 transgene copies per cell. Copy number was assessed as previously described. Each point represents two technical replicates, with error bars showing SEM. Measurements prior to day 17 were saturated (< 5% negative droplets) and therefore could not be accurately reported.

**Supplementary figure 5: In / out PCR analysis gels for detection of TRAC-CAR20 and CD3ζ-CAR20 insertions.**

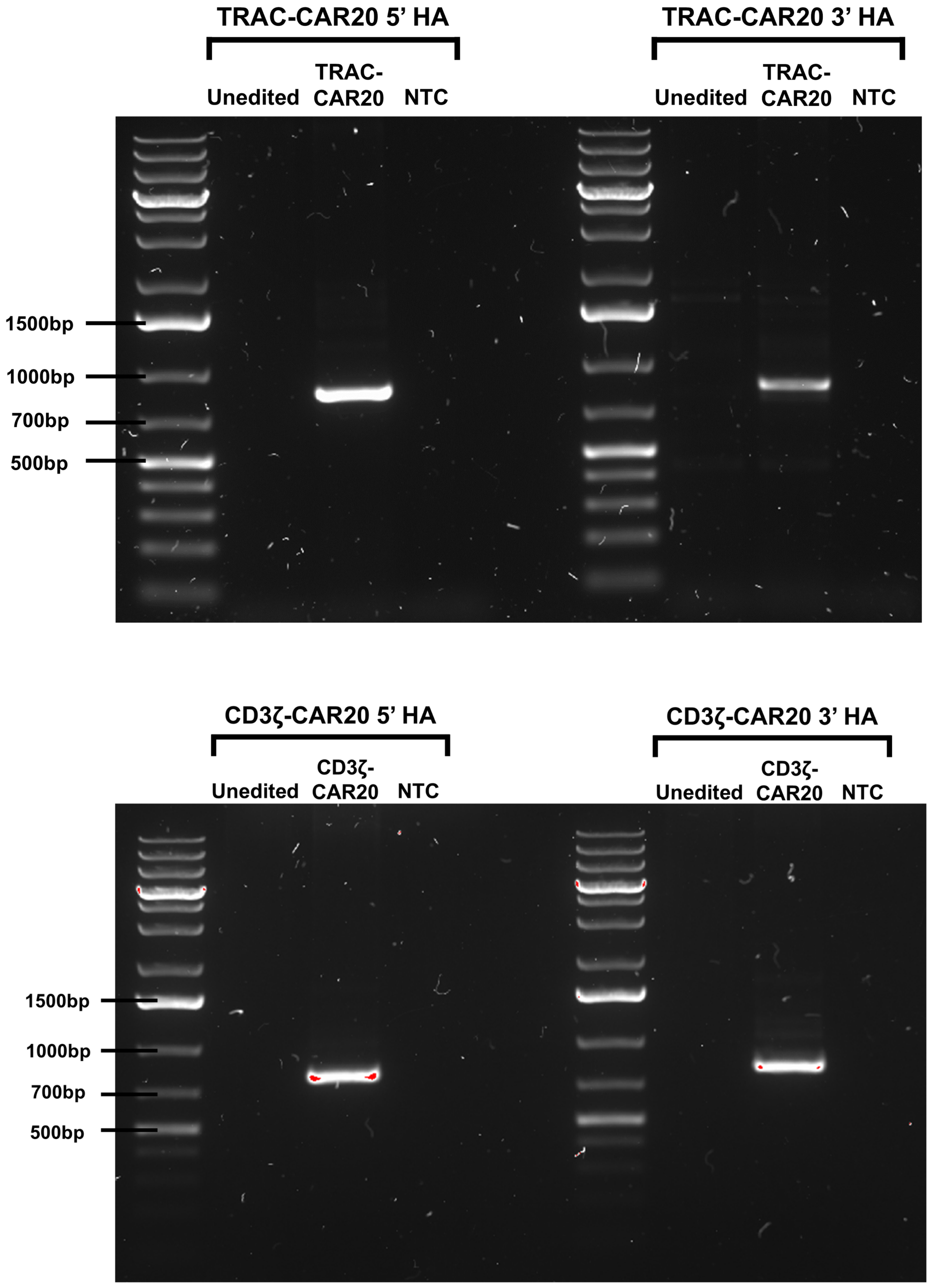


Agarose gels showing PCRs detecting targeted insertion of TRAC-CAR20 and CD3ζ-CAR20. 'In / out' PCR primer pairs were designed at the 5' and 3' ends of each transgene such that they only amplify upon successful insertion of the transgene. Expected band sizes: TRAC-CAR20 5' HA, 832bp; TRAC-CAR20 3' HA, 828bp; CD3ζ-CAR20 5' HA, 769bp; CD3ζ-CAR20 3' HA, 783bp.

### Supplementary figure 6: Additional nanopore analysis figures

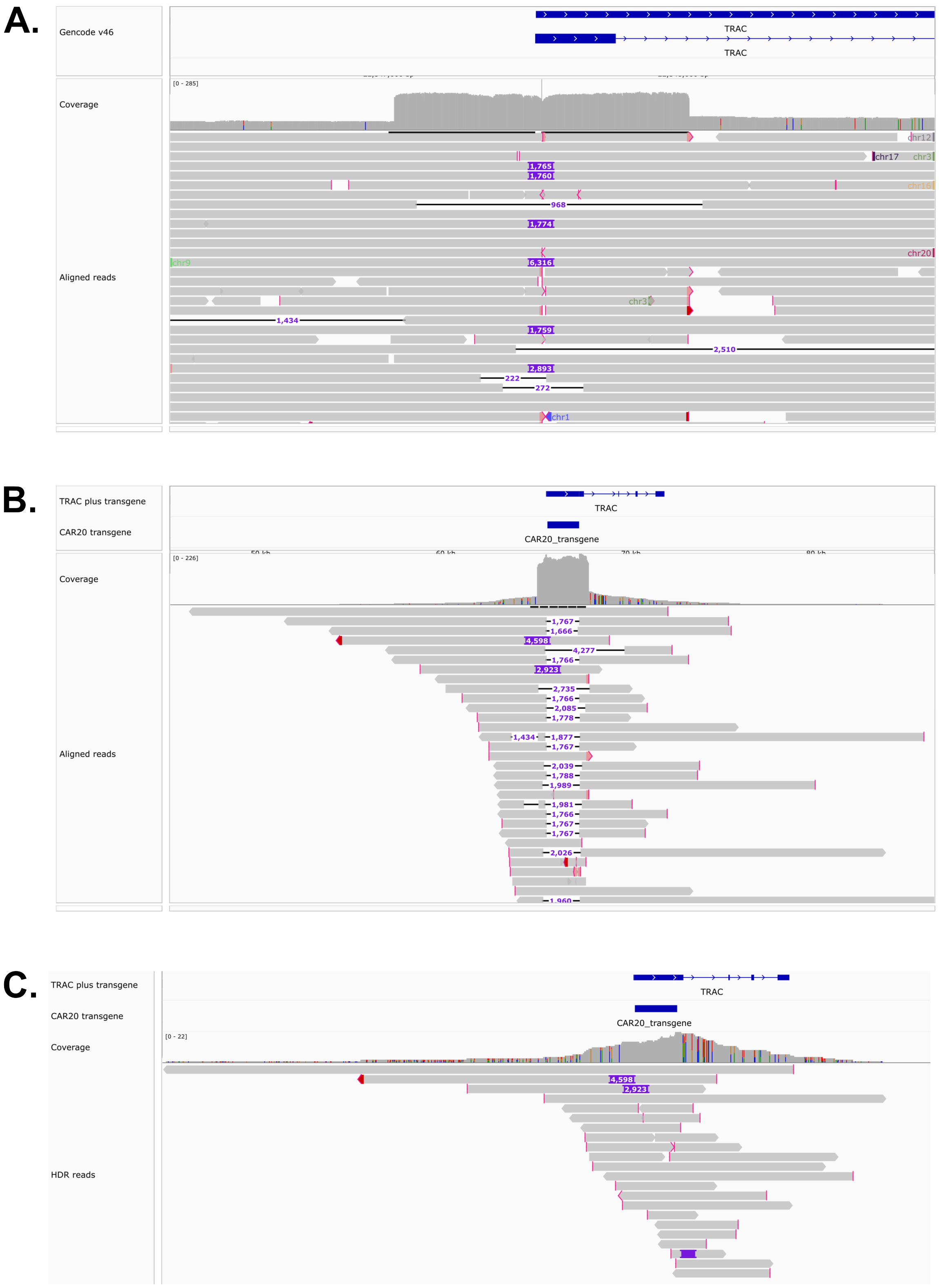

**Supplementary figure 6 (cont.):**

**A.** IGV image showing alignment of TRAC-CAR20 nanopore sequencing reads at *TRAC* exon 1. As can be seen, there is sharp increase in coverage across the 1Kb region corresponding to the 500bp 5' and 3' homology arms of the TRAC-CAR20 donor template. This data suggests that there is still non-integrated donor template present in the end-of-production TRAC-CAR20 cells.

**B.** IGV figure showing alignment of the nanopore sequencing data to a custom reference containing the CAR20 transgene inserted into the *TRAC* locus, prior to any filtering of episomal template reads. As can be seen, there is a substantial spike in coverage across the transgene + homology arm region, resulting from the episomal reads at this site.

**C.** IGV image showing reads classified as either "HDR" or "Concatemeric" after analysis. All reads had coverage both in the transgene and in the genome at the TRAC site outside of the homology arms of the donor template, suggesting they originated from transgene insertions.

**Supplementary figure 7: CD4 / CD8 expression in the CAR20$^+$ subpopulation at end-of-production**. Proportion of cells expressing CD4 and CD8 in the CD2$^+$CAR20$^+$ subpopulation of CAR-T cell batches at end-of-production.

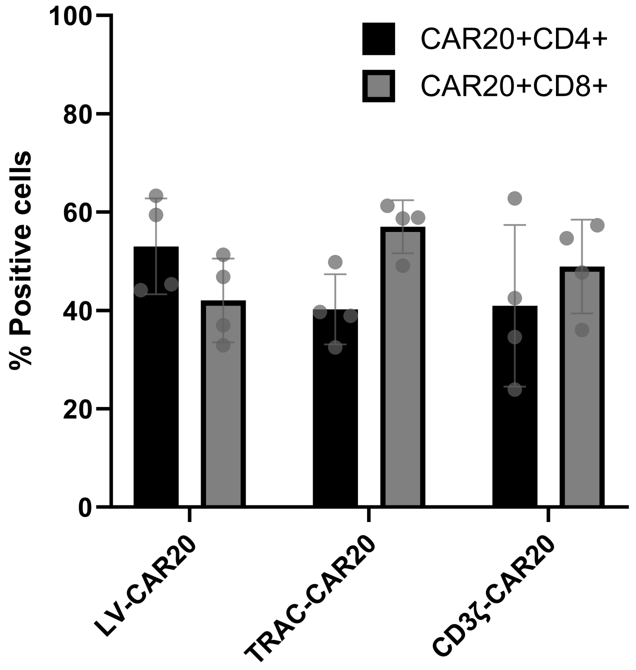

**Supplementary figure 8: Additional donor replicates for $^{51}Cr$ cytotoxicity assays.**

Graphs showing killing of 51Cr-labelled Daudi cells for two additional MNC donors. Similarly to the representative plot in figure 2E, comparable killing was observed for all CAR20-T cell groups.

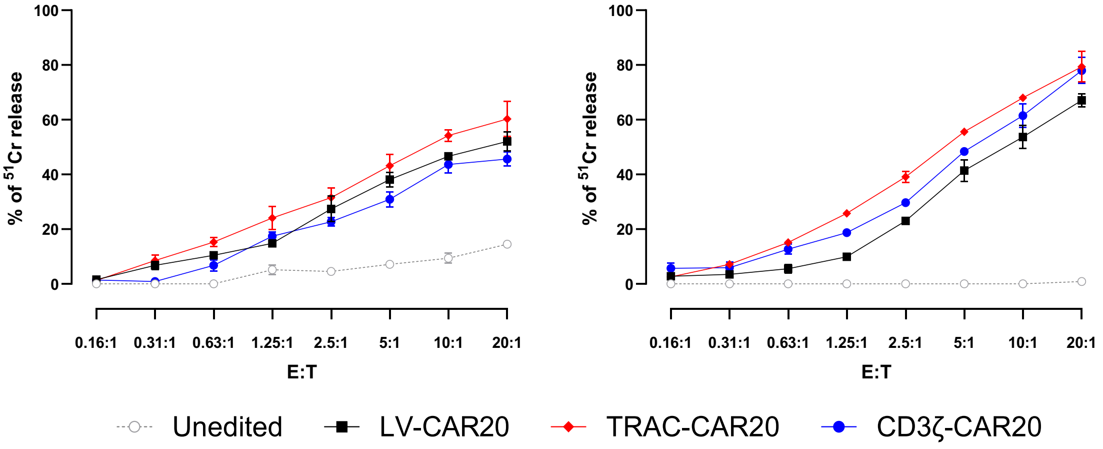

**Supplementary figure 9: Additional human murine chimeric data comparing LV-CAR20, TRAC-CAR20 and CD3ζ-CAR20.**

Graph showing mean radiance values from bioluminescent imaging of Daudi-FLG tumour progression in NSG mouse models from the initial pilot mouse experiment (PBS control, n = 4; for all other groups n = 7). Statistical analysis was carried out by one-way ANOVA of the area-under-the-curve (AUC) for each group, followed by a post-hoc Tukey's test for individual comparisons (with correction for multiple testing). P value summaries: p≥0.05 (*ns*), p<0.05 (*), p<0.01 (**), p<0.001 (***), p≤0.0001 (****).

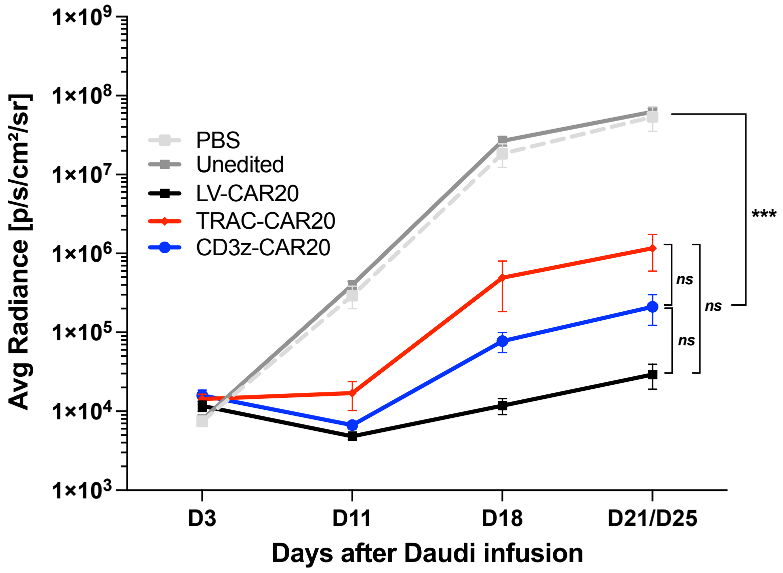

**Supplementary figure 10: Additional phenotyping of CAR-T cells recovered from main *in vivo* comparison of LV-CAR20, TRAC-CAR20 and CAR20-CD3ζ.**

**A.** Graph showing expression of TCRαβ and CAR20 in $CD2^+$ cells recovered from the bone marrow of mice at point of necroscopy. For all treatment groups, the remaining cells were mainly $CAR^+$.

**B.** Graph showing the percentage of cells expressing CD4 and CD8 in the $CD2^+CAR20^+$ sub-population.

A.

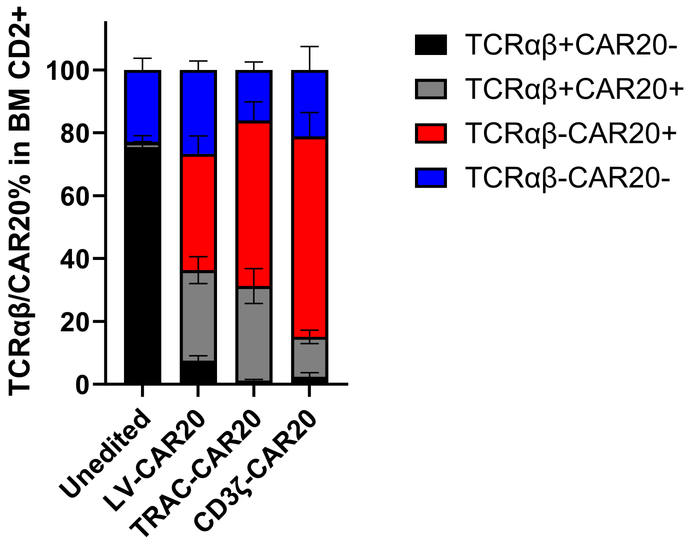


B.

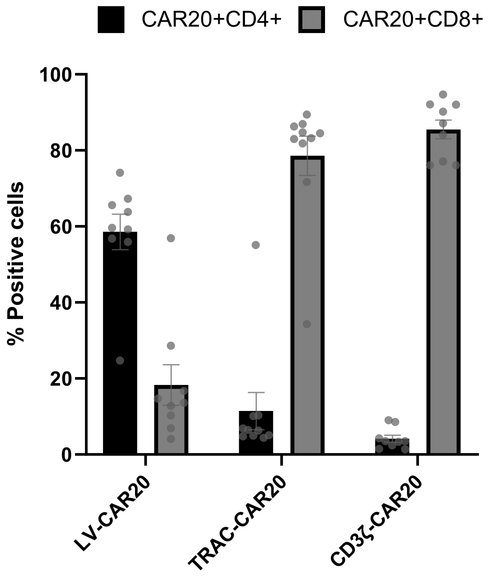

**Supplementary figure 11: Single versus competitive LV-CAR20 / TRAC-CAR20 infusion.**

**A.**

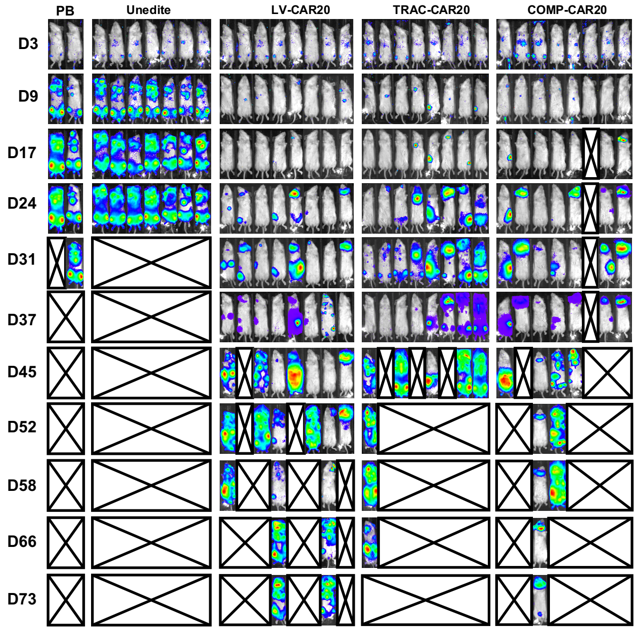


**B.**

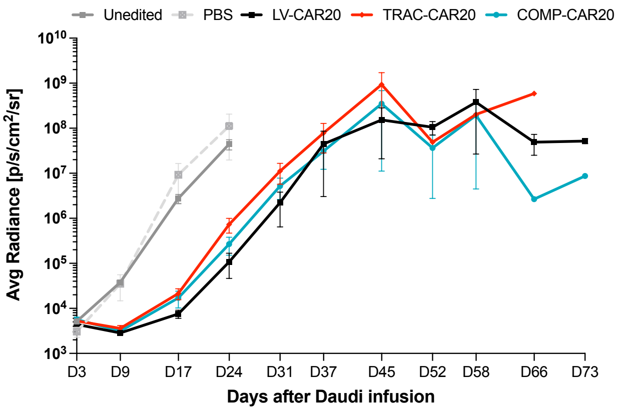


| | D24 cut-off | End of study |
|---|---|---|
| Unedited vs. ALL | *ns* | *NA* |
| LV-CAR20 vs. TRAC-CAR20 | *ns* | *ns* |
| LV-CAR20 vs. COMP-CAR20 | *ns* | *ns* |
| TRAC-CAR20 vs. COMP-CAR20 | *ns* | *ns* |

**C.**

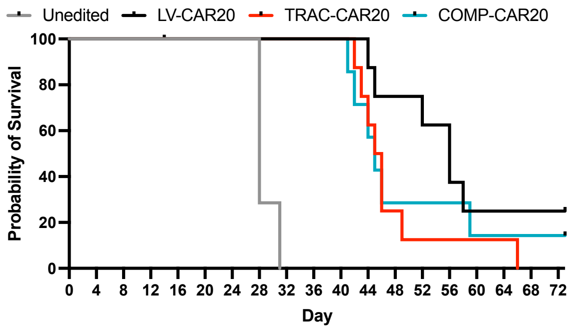


| | P value (*significance*) |
|---|---|
| Unedited vs. LV-CAR20 / TRAC-CAR20 | P < 0.0001 (****) |
| Unedited vs. COMP-CAR20 | P = 0.0002 (***) |
| LV-CAR20 vs. TRAC-CAR20 | P = 0.0517 (*ns*) |
| LV-CAR20 vs. COMP-CAR20 | P = 0.3338 (*ns*) |
| TRAC-CAR20 vs. COMP-CAR20 | P = 0.7781 (*ns*) |

**D.**

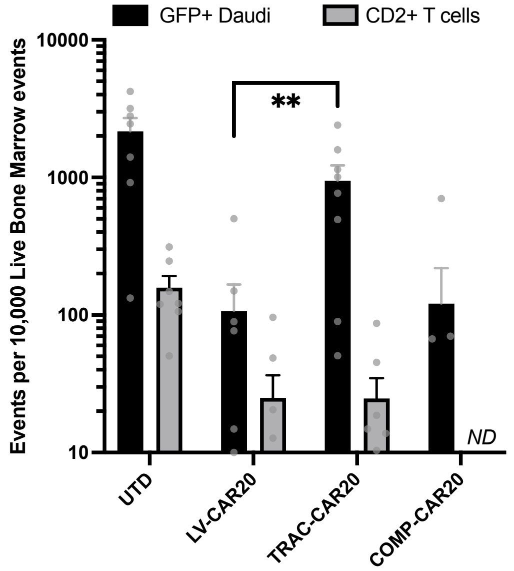


**E.**

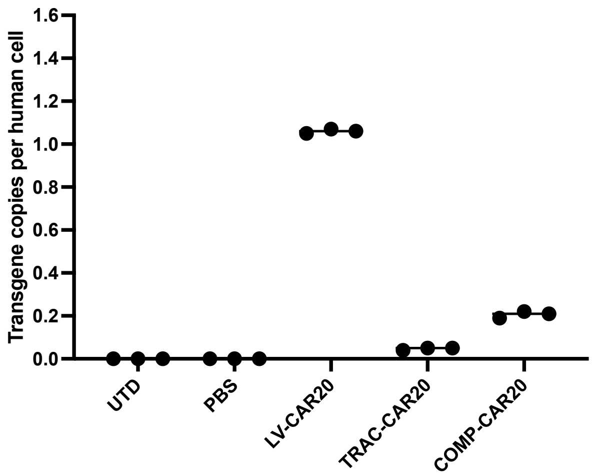


**F.**

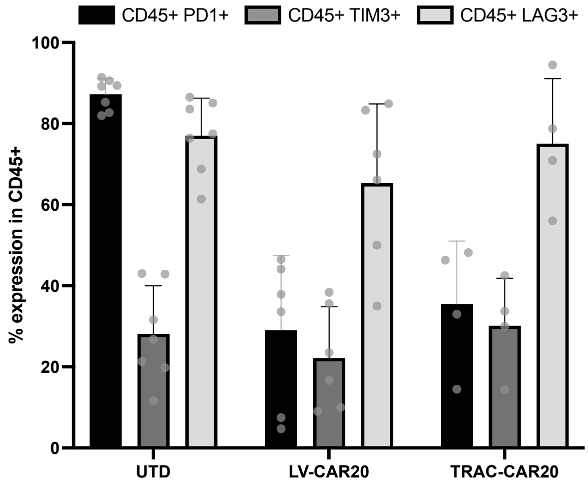

**Supplementary figure 11 (cont.):**

Assessment of control CAR20$^{+}$TCRαβ$^{-}$ cells (2.5x10$^{6}$ CAR$^{+}$ cells i.v.) against Daudi-FLG cells (CD20$^{+}$) in NOD/SCID/γc$^{-/-}$ (NSG) mice.

**A.** Bioluminescent imaging (BLI) of NSG mice engrafted with Daudi-FLG cells and dosed with either LV-CAR20, TRAC-CAR20 or a 1:1 competitive co-infusion of both products (COMP-CAR20). Animals were followed for up to 73 days after tumour engraftment. Mice receiving non-modified T cells or PBS alone provided controls.

**B.** Mean radiance values from BLI of Daudi-FLG leukaemia progression in NSG murine models (PBS control, n = 2; Unedited control, n = 7; for all other groups n = 8). No significant differences were observed in tumour control between the groups. Statistical analysis was carried out by one-way ANOVA of the area-under-the-curve (AUC) for each group, followed by a post-hoc Tukey's test for individual comparisons.

**C.** Overall survival was not significantly different between treatment groups when Kaplan-Meier curves were compared (pairwise log-rank tests to a Bonferroni-corrected threshold of $p<0.05$).

**D.** Daudi-FLG cell counts and CD2$^{+}$ T cell counts in bone marrow (BM) at necroscopy. TRAC-CAR20 animals showed significantly higher mean tumour counts than LV-CAR20 cells at necroscopy (Mann-Whitney U test). Cells were measured by flow cytometry normalised to events per 1x10$^{5}$ live cell events (mean with SEM) (Unedited and COMP-CAR20, n = 7; All other groups n = 8). CD2$^{+}$ T cell counts for COMP-CAR20 were not determined (*ND*).

**E.** ddPCR detection of CAR20 transgene in pooled mouse BM samples taken at necroscopy. Transgene levels detectable in LV-CAR20 animals were elevated compared to other groups. Graph shows the mean of 3 technical replicates per group.

**F.** Expression of PD-1, TIM3 and LAG3 in CD45$^{+}$ cells recovered from the bone marrow showed no significant differences (Unedited, n = 7; LV-CAR20, n = 6; TRAC-CAR20, n = 4).

P value summaries: $p\geq0.05$ (ns), $p<0.05$ (*), $p<0.01$ (**), $p<0.001$ (***), $p\leq0.0001$ (****).

**Supplementary table 1: PCR primers and ddPCR probes**

| Primer set | Oligo type | Sequence |
|---|---|---|
| *TRAC* ICE amplicon | Primer_F | TTGATAGCTTGTGCCTGTCCC |
| | Primer_R | GGCAAACAGTCTGAGCAAAGG |
| *CD3ζ* ICE amplicon | Primer_F | AGGTTTTCCAGGAGCTGGTTG |
| | Primer_R | TCTTCCAATCTCTGCCCACAC |
| *TRAC* 5' in/out | Primer_F | GGCTTAGACGCAGGTGTTCT |
| | Primer_R | GCTTGACCCAGTGCATGTTG |
| *TRAC* 3' in/out | Primer_F | ACACCTATGATGCCCTGCAC |
| | Primer_R | AGCAGTAAGGGGCAAACAGT |
| *CD3ζ* 5' in/out | Primer_F | TTGTCTCCCTGTGGGTAGGC |
| | Primer_R | GCTTGACCCAGTGCATGTTG |
| *CD3ζ* 3' in/out | Primer_F | CCGTGCATACCAGAGGACTG |
| | Primer_R | TGGGTTGAGAAGCACTGAAGT |
| TRAC-CAR20 PCR template production | Primer_F | TCAGGTTTCCTTGAGTGGCA |
| | Primer_R | CATTCCTGAAGCAAGGAAACAG |
| CD3z-CAR20 PCR template production | Primer_F | GCCACATCTGCCGTTGGTGC |
| | Primer_R | ACCAGTAGCATCGCCTTCCC |
| CAR20 scFv ddPCR assay | Primer_F | TTCACCAGCTACAACATGC |
| | Primer_R | TAGTACACGGCGCTATCTT |
| | Probe_FAM | ACGGCGACACCTCCTACAAC |
| Human albumin ddPCR assay | Primer_F | GCTGCTATCTCTTGTGGGCTGT |
| | Primer_R | ACTCATGGGAGCTGCTGGTTC |
| | Probe_HEX | CCTGTCATGCCCACACAAATCTCTCC |

**Supplementary table 2: sgRNA sequences**

| sgRNA | Sequence |
|---|---|
| *TRAC* KO/KI | TCTCTCAGCTGGTACACGGC |
| *CD3ζ* KO/KI | CACCTTCACTCTCAGGAACA |
| *TRAC* double-tap 1 | AGTCTCTCAGCTGGTACGGC |
| *TRAC* double-tap 2 | GTCTCTCAGCTGGTACAGGC |
| *TRAC* double-tap 3 | CTCTCAGCTGGTACACCGGC |
| *B2M* | ACTCACGCTGGATAGCCTCC |
| *CD38* | GCGCCAGCAGTGGAGCGGTC |

**Supplementary table 3: Key antibodies used in study**

| Target (all human) | Conjugate | Clone | Supplier | Product reference |
|---|---|---|---|---|
| CD2 | VioBlue | LT2 | Miltenyi Biotec | 130-098-701 |
| TCRαβ | PE | BW242/412 | Miltenyi Biotec | 130-113-531 |
| TCRαβ | Biotin | BW242/412 | Miltenyi Biotec | 130-113-529 |
| CD4 | PE-Vio770 | REA623 | Miltenyi Biotec | 130-113-227 |
| CD8 | APC-Vio770 | REA734 | Miltenyi Biotec | 130-110-681 |
| Rituximab | FITC | MB2A4 | BioRad | MCA2260F |
| PD-1 | PE | PD1.3.1.3 | Miltenyi Biotec | 130-117-384 |
| PD-1 | BB515 | EH12.1 | BD Biosciences | 565936 |
| TIM-3 | AF647 | 7D3 | BD Biosciences | 565559 |
| LAG-3 | BV421 | T47-530 | BD Biosciences | 565720 |
| CD62L | APC | 145/15 | Miltenyi Biotec | 130-113-617 |
| CD45RA | PE-Vio770 | REA1047 | Miltenyi Biotec | 130-117-746 |
